# Synthesis of K^+^ channel radioligand [^18^F]5-methyl-3-fluoro-4-aminopyridine and PET imaging in mice

**DOI:** 10.1101/2024.07.19.604281

**Authors:** Yang Sun, Karla M. Ramos-Torres, Kazue Takahashi, Lauren L. Zhang, Pedro Brugarolas

## Abstract

[^18^F]3-fluoro-4-aminopyridine ([^18^F]3F4AP) is the first positron emission tomography (PET) radioligand that targets voltage-gated potassium (K^+^) channels in the brain for imaging demyelination. [^18^F]3F4AP exhibits high brain penetration, favorable kinetics for PET imaging, and high sensitivity to demyelinating lesions. However, recent studies in awake human subjects indicate lower metabolic stability than in anesthetized animals, resulting in reduced brain uptake. Therefore, there is a need for novel radioligands for K^+^ channels with suitable pharmacological properties and enhanced metabolic stability. Recent *in vitro* studies demonstrate that 5-methyl-3-fluoro-4-aminopyridine (5Me3F4AP) exhibits comparable binding affinity to K^+^ channels, p*K*_a_, logD, and membrane permeability as 3F4AP, and a slower enzymatic metabolic rate, suggesting its potential as a K^+^ channel PET tracer. In this study, we describe the radiochemical synthesis of [^18^F]5Me3F4AP using an isotope exchange method from the corresponding 3-fluoro-5-methyl-4-nitropyridine N-oxide, followed by a palladium on carbon mediated hydrogenation of the nitro and N-oxide groups. This method yielded [^18^F]5Me3F4AP with high purity and acceptable molar activity. PET/CT studies using naïve mice demonstrate that [^18^F]5Me3F4AP effectively crosses the blood-brain barrier and has comparable kinetics to [^18^F]3F4AP. These findings strongly suggest that [^18^F]5Me3F4AP is a promising candidate for neuroimaging applications and warrant further studies to investigate its sensitivity to lesions and *in vivo* metabolic stability.

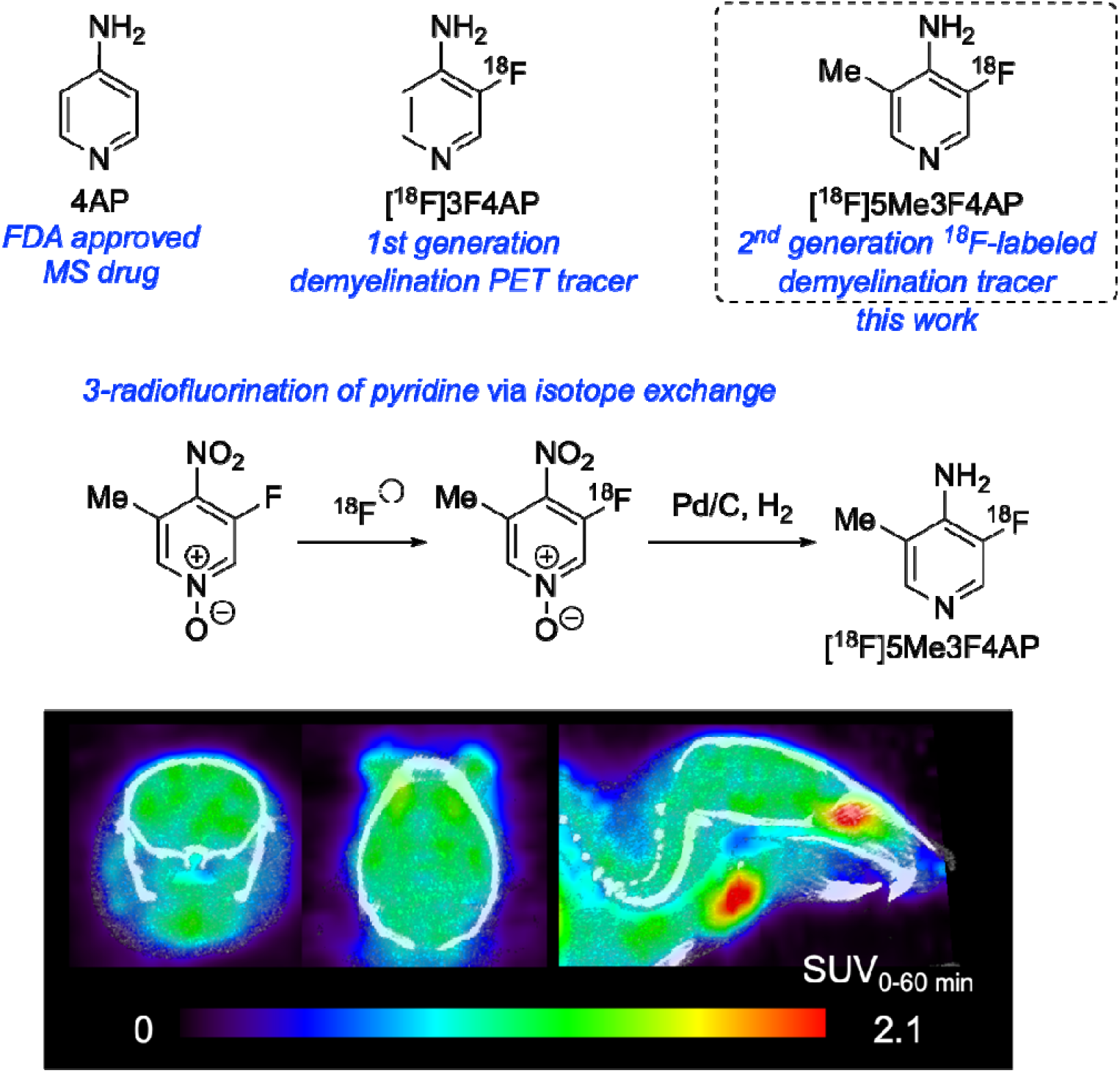

## Introduction

4-Aminopyridine (4AP) is a potassium (K^+^) channel blocker approved by the FDA for improving walking speed in people with multiple sclerosis (MS)^1^. By binding to and blocking K^+^ channels in demyelinated axons, 4AP enhances nerve conduction^2-5^. Based on this compound, the first-generation positron emission tomography (PET) radioligand, [^18^F]3-fluoro-4-aminopyridine ([^18^F]3F4AP), was developed^6, 7^. This PET tracer showed high binding to demyelinated lesions in rodent models of MS^6^, a rat model of spinal cord injury^8^, and a monkey with a traumatic brain injury^9^. Given the promising results in animal models, this compound has recently been advanced to human research studies (Clinicaltrials.gov NCT04699747). Studies in healthy volunteers showed good brain penetration, fast kinetics and low radiation dosimetry^10^ but, also, lower metabolic stability than what had been observed in animal studies. Additional studies into the metabolism of [^18^F]3F4AP indicated that in the absence of isoflurane anesthetics that can inhibit the metabolic enzyme CYP2E1, [^18^F]3F4AP primarily underwent oxidation at the 5-position^11, 12^. In parallel, another carbon-11 labeled radioligand, [^11^C]3-methyl-4-aminopyridine ([^11^C]3Me4AP), demonstrated higher *in vitro* and *in vivo* binding affinity to K^+^ channels but exhibited slower kinetics due to its lower brain penetration^13^.

Given the favorable half-life of ^18^F (110 min), we aimed to identify an ^18^F-labeled tracer with superior *in vivo* properties compared to [^18^F]3F4AP. We recently investigated the chemical and biophysical properties of 5-methyl-3-fluoro-4-aminopyridine (5Me3F4AP) through several *in vitro* experiments^14^. This trisubstituted compound was designed to combine the advantageous attributes of [^18^F]3F4AP and [^11^C]3Me4AP while blocking the reactive 5-position. 5Me3F4AP demonstrated a binding affinity to K^+^ channels similar to that of 3F4AP and 4AP, with relative IC_50_ values to 4AP being 1 for 5Me3F4AP and 0.8 for 3F4AP (**Table 1**). This makes it the first trisubstituted 4AP derivative reported to have high binding affinity. The p*K*_a_ of this compound, close to physiological pH, and its positive logD (0.66, indicating higher lipophilicity), were comparable to 3F4AP but distinct from 4AP and 3Me4AP, suggesting enhanced brain penetration and favorable kinetics. This is because only the neutral form of the molecule, with good lipophilicity, can effectively cross the lipid-rich blood-brain barrier (BBB). In an artificial membrane permeability test, 5Me3F4AP exhibited a permeability rate three times faster than 3F4AP and 18 times faster than 4AP, consistent with expectations based on their p*K*_a_ and logD values. Furthermore, 5Me3F4AP demonstrated increased resistance to CYP2E1 enzymatic metabolic degradation. An *in vitro* enzyme assay using CYP2E1, a critical enzyme for metabolizing 4AP derivatives, revealed that 5Me3F4AP is metabolized at half the rate of 3F4AP.

**Table 1.**
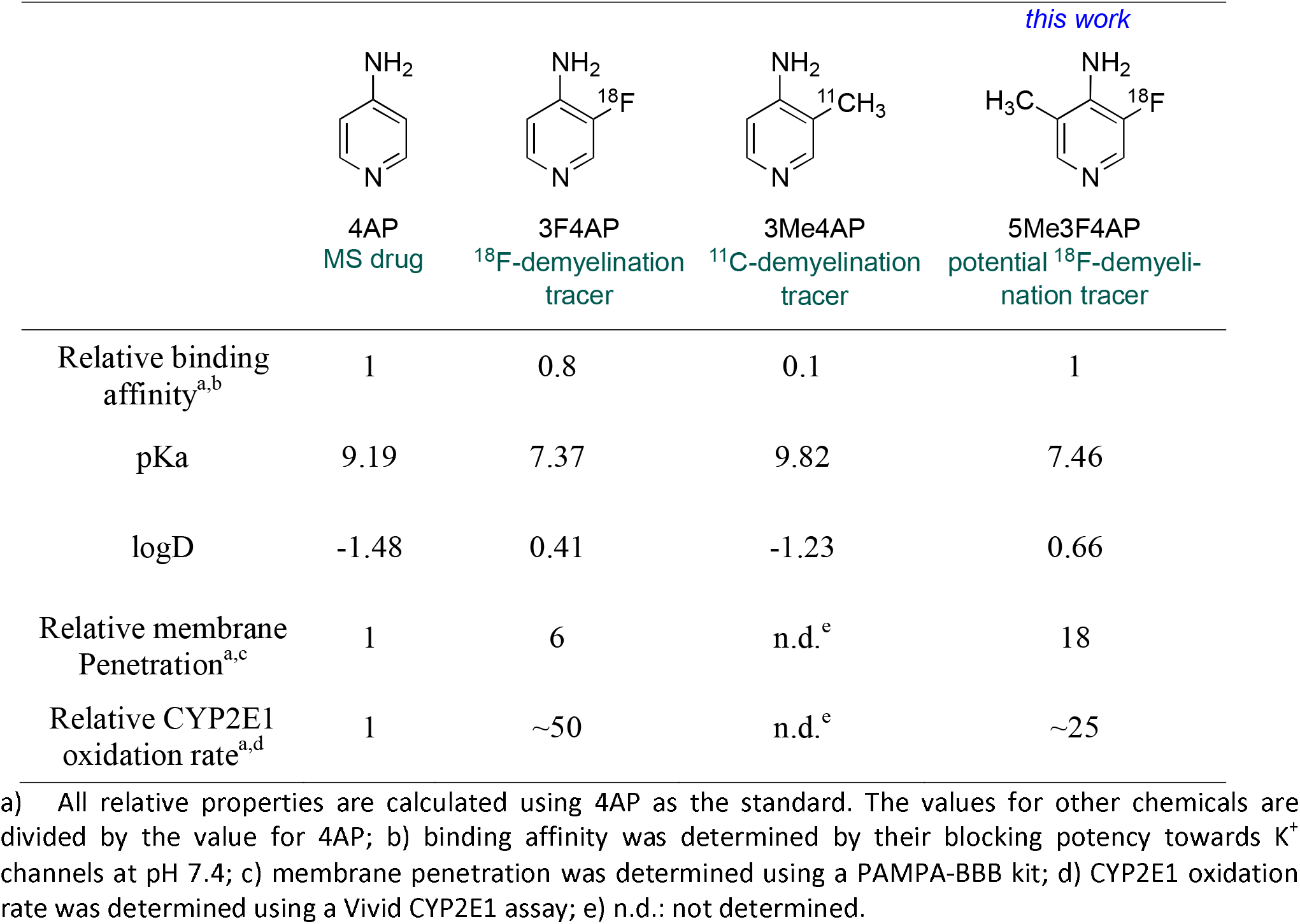
In vitro properties comparison between four 4AP derivatives.

|  |  |  |  | *this work* |
| --- | --- | --- | --- | --- |
|  | 4AP<br>MS drug | 3F4AP<br><sup>18</sup>F-demyelination<br>tracer | 3Me4AP<br><sup>11</sup>C-demyelination<br>tracer | 5Me3F4AP<br>potential <sup>18</sup>F-demyeli-<br>nation tracer |
| Relative binding<br>affinity<sup>a,b</sup> | 1 | 0.8 | 0.1 | 1 |
| pKa | 9.19 | 7.37 | 9.82 | 7.46 |
| logD | -1.48 | 0.41 | -1.23 | 0.66 |
| Relative membrane<br>Penetration<sup>a,c</sup> | 1 | 6 | n.d.<sup>e</sup> | 18 |
| Relative CYP2E1<br>oxidation rate<sup>a,d</sup> | 1 | ~50 | n.d.<sup>e</sup> | ~25 |

a) All relative properties are calculated using 4AP as the standard. The values for other chemicals are divided by the value for 4AP; b) binding affinity was determined by their blocking potency towards K<sup>+</sup> channels at pH 7.4; c) membrane penetration was determined using a PAMPA-BBB kit; d) CYP2E1 oxidation rate was determined using a Vivid CYP2E1 assay; e) n.d.: not determined.

These compelling findings prompted us to evaluate the imaging properties of 5Me3F4AP *in vivo*. Additionally, we aimed to investigate effective methods for the radiofluorination of this compound, given the well-documented challenges associated with fluorinating pyridines at the 3- or 5-positions. Typical approaches include nucleophilic aromatic substitution (SNAr) reactions of precursors containing nitro group^15^, iodonium salt/ylide^16-18^, sulfonium salt^19^, N-sydnone^20^, or halide^21^, and copper-mediated radiofluorination of pyridinyl boronic acid/ester precursors^22-24^. One significant challenge is the low yield in reactions without electron-withdrawing substituents or in copper-mediated reactions^24^. Additionally, the accessibility of some precursors, such as iodonium salts or ylides, is limited due to the oxidation conditions required to form the iodonium salt/ylide, which often leads to the oxidation of the pyridine ring, especially in cases lacking strong electron-withdrawing groups^18^. Furthermore, yields dramatically decrease with bulkier para-substitutions^16, 17^. In this work, we investigate an isotope exchange approach for synthesizing [^18^F]5Me3F4AP, aiming to overcome these synthetic hurdles and achieve a viable radiolabeled product for neuroimaging applications.

## Results and discussion

### Radiochemical Synthesis of [^18^F]5Me3F4AP

Initial attempts to synthesize [^18^F]5Me3F4AP focused on copper-mediated radiofluorination of a 3-boronic ester precursor (**1**), a widely used method for aromatic radiofluorination^22-28^. This was to be followed by the reduction of the 4-nitro group to yield the desired [^18^F]5Me3F4AP (**Figure 1A**). Despite its popularity and success in para-radiofluorination of pyridine^24^, initial attempts to radiofluorinate this Bpin precursor using standard conditions did not yield the desired meta-radiofluorinated product and this approach was abandoned.

**Figure 1.**
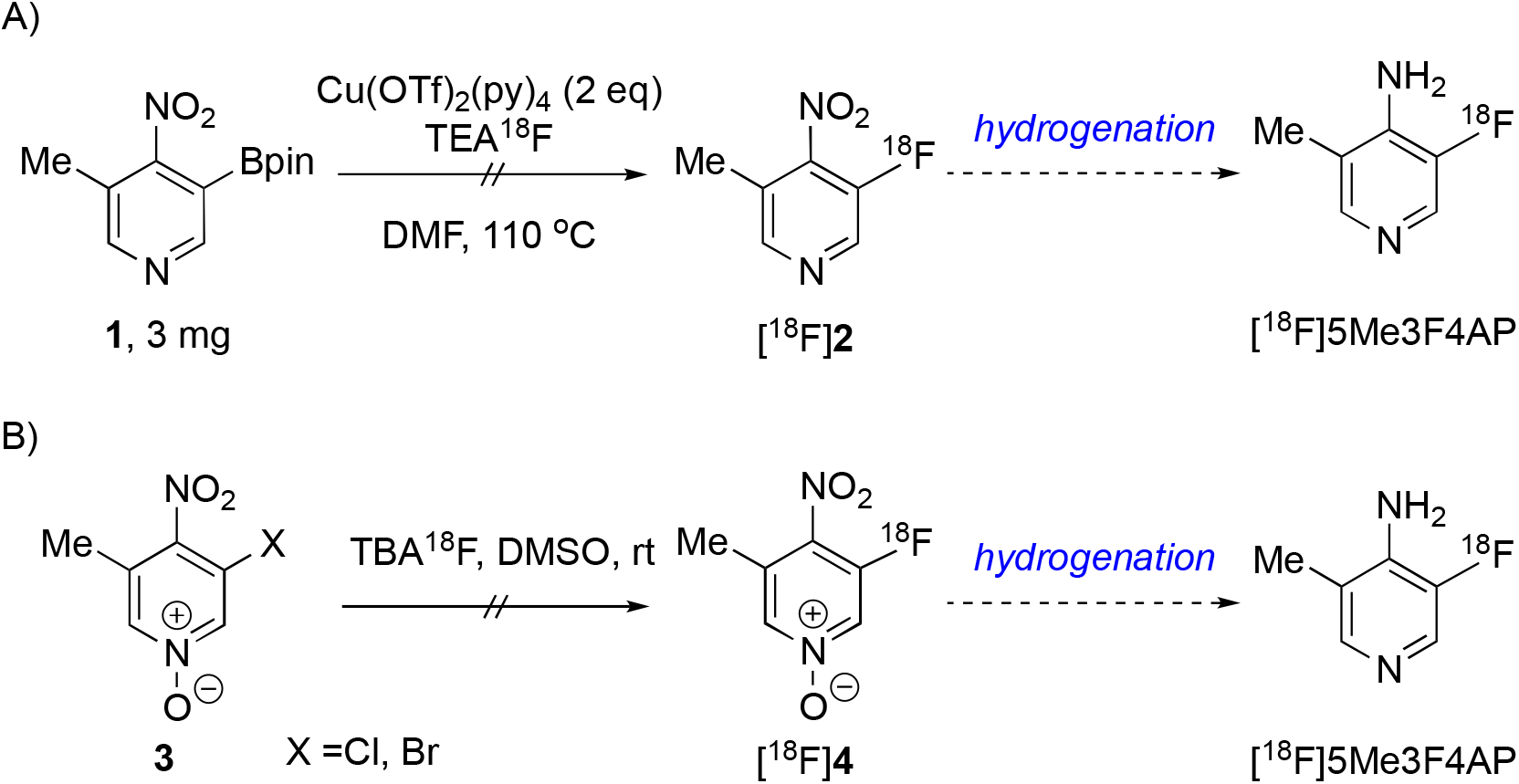
Two proposed routes for radiosynthesis of [^18^F]5Me3F4AP.

We then explored an alternative halogen exchange method using the corresponding 3-chloro/bromo-5-methyl-4-nitro pyridine N-oxide precursor (**3**), followed by a one-step hydrogenation to reduce the 4-nitro group and N-oxide (**Figure 1B**) paralleling previously successful in the synthesis of [^18^F]3F4AP^29^. In the case of 3F4AP performing the reaction at room temperature resulted in halogen exchange at the 3-position from the Br reagent in 10.4±1.8 % decay-corrected isolated radiochemical yield (RCY) whereas reactions conducted at high temperatures predominantly resulted in denitrofluorination at the 4-position. However, during the synthesis of [^18^F]5Me3F4AP, we encountered poor stability of the precursor in the presence of a base even at room temperature, which resulted in a RCY of less than 5%.

In addition to testing the chloro- and bromo-precursors, we also tested the corresponding fluoro-precursor *via* isotopic exchange (**Table 2**). Manual labeling with aliquoted azeotropic [^18^F]fluoride ([^18^F]F^-^) solution in various solvents at room temperature for 15 minutes showed that non-protic polar solvents such as DMSO, MeCN, and THF yielded moderate RCYs with tetrabutylammonium [^18^F]fluoride (entries 1-4). However, no product was obtained with potassium [^18^F]KF and kryptofix 222 (K222) (entry 5). The low RCY was typically accompanied by the observation of precursor decomposition. This may be because KF/K222 is less nucleophilic and more basic than [^18^F]TBAF, leading to less reactivity and more severe precursor decomposition. When the labeling process was transferred to an automated synthesizer, the increased base ratio significantly decreased the RCY due to precursor decomposition caused by the higher amount of base required for efficient elution of the [^18^F]F^-^ from the QMA cartridge (entry 6, elution efficiency = 71 ± 9%). Therefore, a shorter reaction time was tested to minimize decomposition. This change led to a drastic improvement in RCY (entry 7, representative traces in **Fig. 2C**). This also confirms that the ^18^F/^19^F exchange occurs almost immediately. Using a minimal amount of precursor, which showed no significant reduction in conversion (entries 8 *vs*. 7) and reacting for only 1 minute resulted in a 37 ± 9% isolated yield after quenching with the addition of one equivalent of acetic acid (AcOH) to the bicarbonate base (entry 8).

**Table 2.**
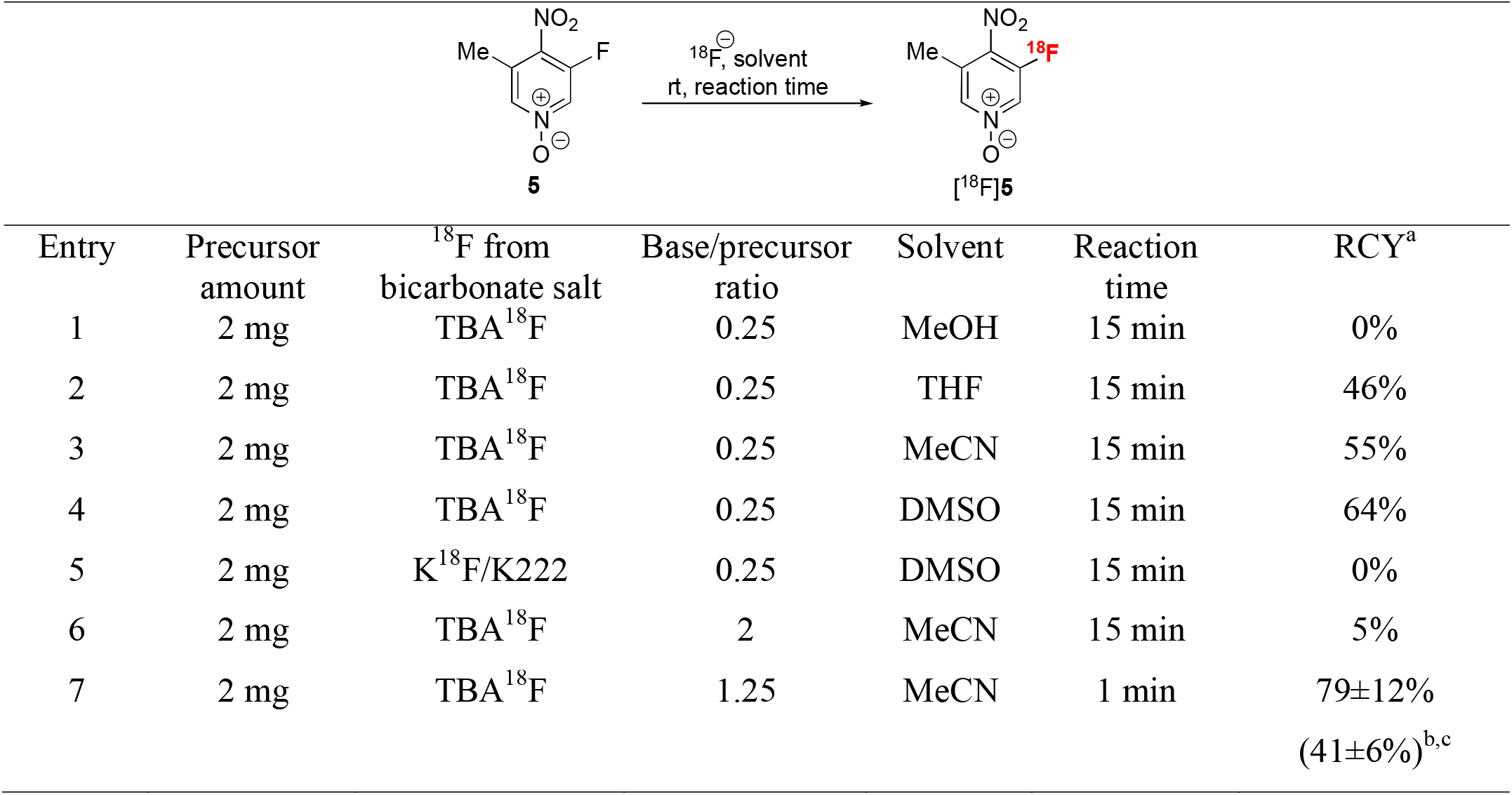
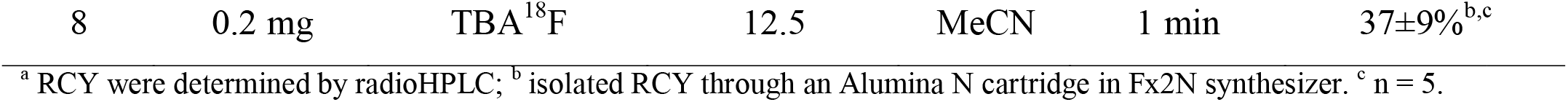
Radiochemical synthesis by ^18^F/^19^F exchange.

|  |  |  |  |  |  |  |
| --- | --- | --- | --- | --- | --- | --- |
| Entry | Precursor amount | $^{18}\text{F}$ from bicarbonate salt | Base/precursor ratio | Solvent | Reaction time | RCY<sup>a</sup> |
| 1 | 2 mg | TBA$^{18}\text{F}$ | 0.25 | MeOH | 15 min | 0% |
| 2 | 2 mg | TBA$^{18}\text{F}$ | 0.25 | THF | 15 min | 46% |
| 3 | 2 mg | TBA$^{18}\text{F}$ | 0.25 | MeCN | 15 min | 55% |
| 4 | 2 mg | TBA$^{18}\text{F}$ | 0.25 | DMSO | 15 min | 64% |
| 5 | 2 mg | K$^{18}\text{F}$/K222 | 0.25 | DMSO | 15 min | 0% |
| 6 | 2 mg | TBA$^{18}\text{F}$ | 2 | MeCN | 15 min | 5% |
| 7 | 2 mg | TBA$^{18}\text{F}$ | 1.25 | MeCN | 1 min | $79 \pm 12\%$<br>$(41 \pm 6\%)^{\text{b,c}}$ |
| 8 | 0.2 mg | TBA<sup>18</sup>F | 12.5 | MeCN | 1 min | 37±9%<sup>b,c</sup> |

<sup>a</sup> RCY were determined by radioHPLC; <sup>b</sup> isolated RCY through an Alumina N cartridge in Fx2N synthesizer. <sup>c</sup> n = 5.

**Figure 2.**
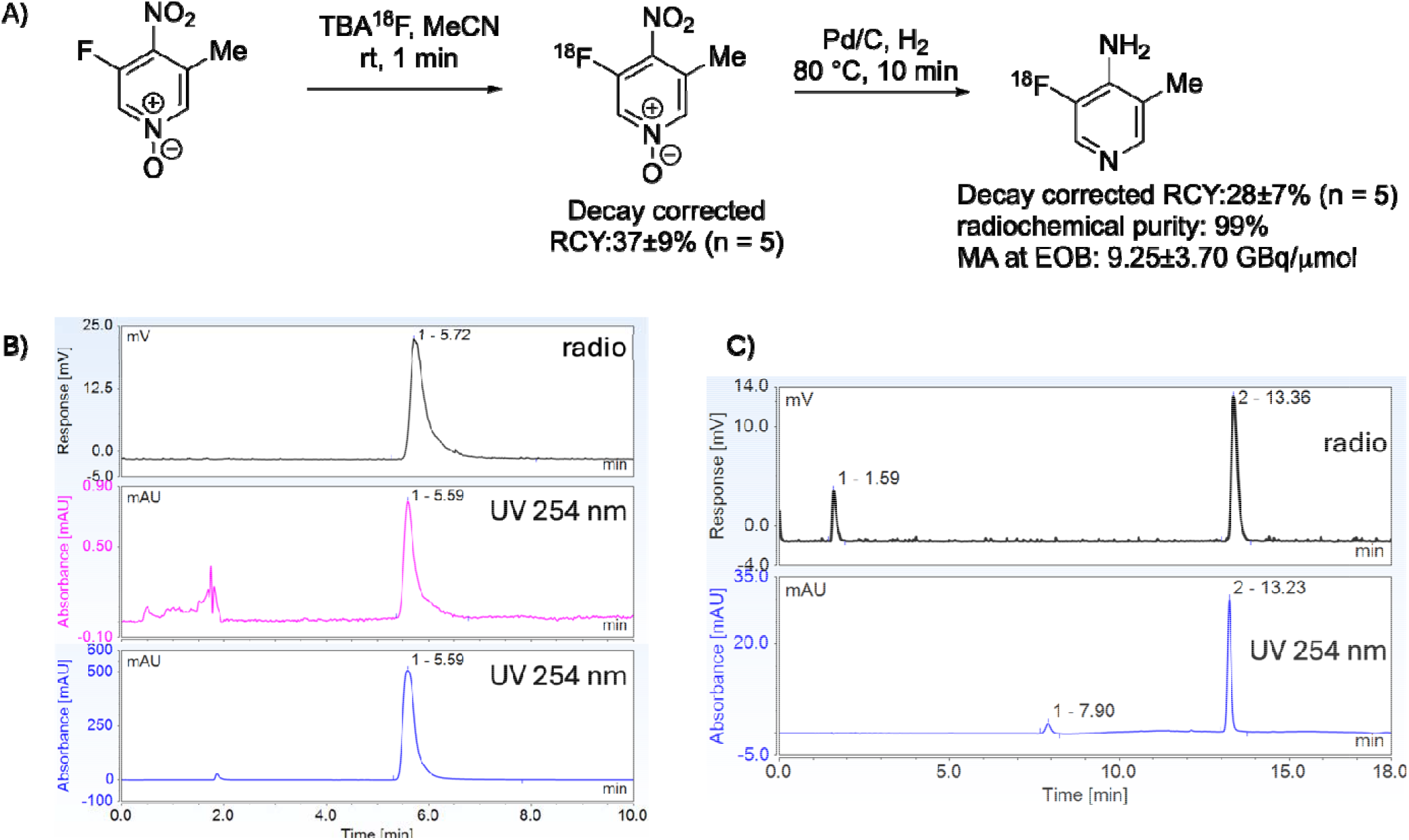
A) Chemical scheme of the radiosynthesis of [^18^F]5Me3F4AP; B) Quality control of [^18^F]5Me3F4 AP (radioactivity trace at the top, UV 254 nm trace in the middle, and the non-radioactive standard at the bottom) ; C) Representative HPLC chromatogram of the reaction crude from step 1 (radioactivity trace at the top and UV 254 nm trace at the bottom).

Subsequently, a palladium on carbon catalyzed hydrogenation was employed to reduce the N-oxide and nitro group in a single step, achieving a 60-90% RCY in 10 min. Starting from 0.2 mg (1.2 µmol) of precursor, we used this rapid isotope exchange labeling reaction, quenched by AcOH, followed by a one-step reduction with palladium on carbon and hydrogen (**Fig. 2A**). Despite some losses due to transfer and filtration of the reaction crude, this method yielded the desired [^18^F]5Me3F4AP with a 28 ± 7% decay corrected RCY (n = 5), >99% radiochemical purity, and a moderate molar activity of 9.25 ± 3.70 GBq/µmol (**Fig. 2B**). While high MA is critical for many brain tracers due to specific binding considerations^30^, this does not seem to be the case for 3F4AP. In studies in rhesus macaques, adding cold 3F4AP to [^18^F]3F4AP doses (effectively lowering the MA) resulted in increased tracer binding in the regions of interest^9^. This phenomenon may be attributed to compensatory mechanism by which blocking a fraction of voltage-gated potassium channels causes more to open.

### PET/CT imaging in naïve mice

After successfully synthesizing [^18^F]5Me3F4AP, we conducted PET/CT studies in naïve mice to evaluate its brain penetration and kinetics. Dynamic PET images were captured from 0 to 60 minutes post-administration of the radioligands, focusing on the whole brain (n = 2 for each tracer). Representative images in **Figure 3B** show the average distribution and accumulation of the tracer within the brain over the 60-minute period, indicating that [^18^F]5Me3F4AP effectively crosses the BBB and binds to its target within the brain. Additionally, high uptake in the eyes and salivary glands was also consistent with [^18^F]3F4AP.

**Figure 3.**
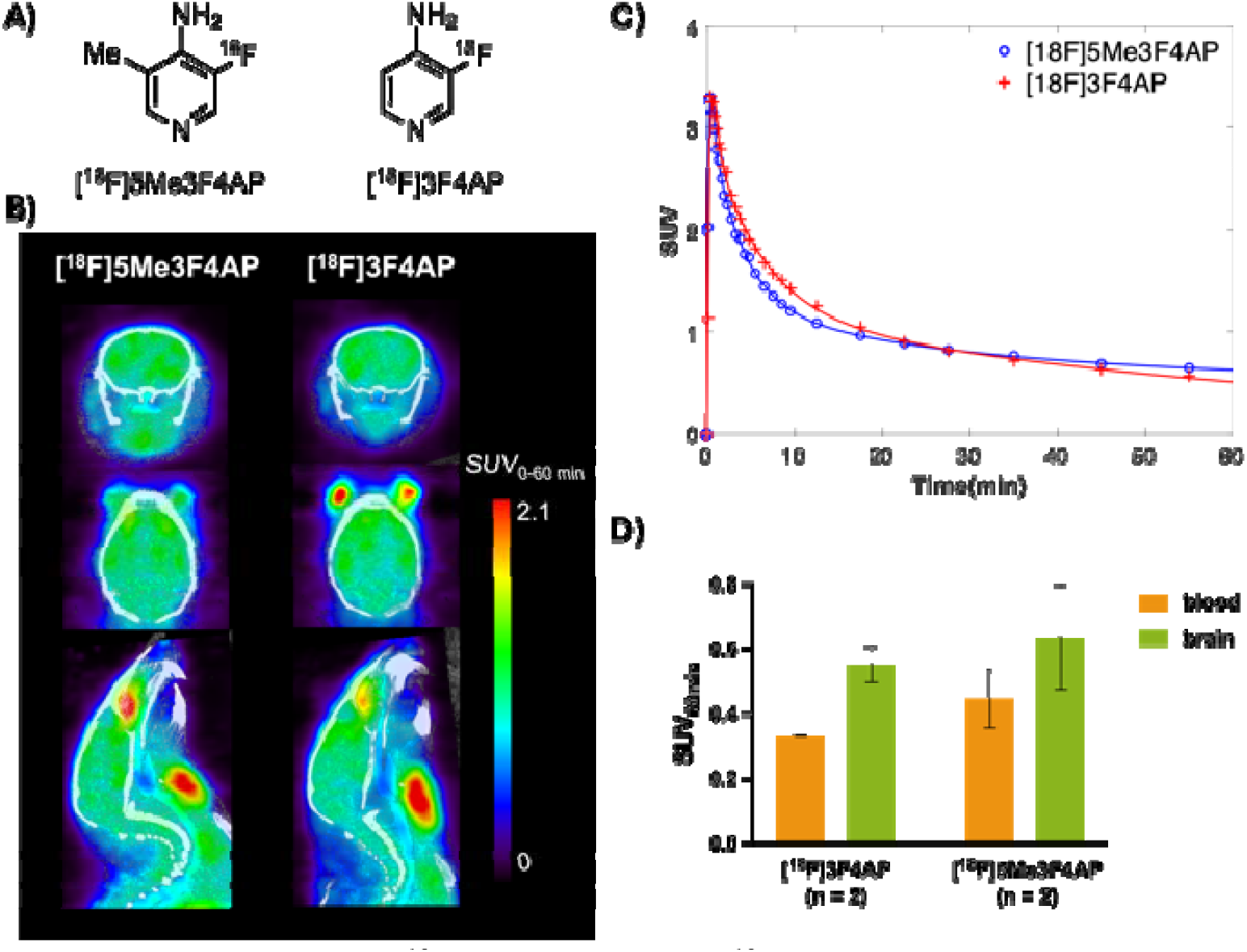
A) Structure of [^18^F]5Me3F4AP and [^18^F]3F4AP; B) Representative summed PET images captured from 0-60 minutes for each tracer; C) Time-activity curves; D) Terminal uptake in blood and brain calculated by *ex vivo* gamma counting.

To quantitatively assess the tracer kinetics, we generated time-activity curves (TACs) for the whole brain (**Fig. 3C**). The reproducibility was also high, with almost identical curves observed in two different animals. The TACs indicated that [^18^F]5Me3F4AP reached a peak standardized uptake value (SUV) of approximately 3.3 within 1 minute, similar to [^18^F]3F4AP. Furthermore, the washout rate of [^18^F]5Me3F4AP was comparable to that of [^18^F]3F4AP, suggesting that both tracers have similar brain penetration, binding, and clearance dynamics. In contrast, [^11^C]3Me4AP did not exhibit such favorable dynamics^13^. Terminal brain uptake was measured by gamma counting post-imaging. Results showed that at the end of the imaging period, the concentrations of [^18^F]5Me3F4AP in both the blood (SUV_[18F]5Me3F4AP_ = 0.45 ± 0.07 vs. SUV_[18F]3F4AP_ = 0.33 ± 0.01) and brain (SUV_[18F]5Me3F4AP_ = 0.64 ± 0.12 vs. SUV_[18F]3F4AP_ = 0.55 ± 0.04) were also comparable to those of [^18^F]3F4AP (**Fig. 3D**) and consistent with the imaging SUV data.

Given that [^18^F]3F4AP demonstrates excellent imaging properties in anesthetized animals, the comparable kinetics and brain uptake of [^18^F]5Me3F4AP hold substantial promise for further investigation in animal models of disease and under awake conditions. Its potential sensitivity to lesions and metabolic stability are crucial factors were [^18^F]5Me3F4AP may demonstrate superiority over [^18^F]3F4AP. These initial results provide a solid foundation for [^18^F]5Me3F4AP as a viable brain imaging tracer, with potential advantages in kinetics, brain penetration and retention.

## Conclusion

Our study underscores the potential of methylated [^18^F]3F4AP as a neuroimaging tracer. Designed for enhanced brain retention and favorable kinetics, it achieved promising results in initial PET/CT studies. Despite challenges in radiochemical synthesis, our isotope exchange approach yielded [^18^F]5Me3F4AP with satisfactory purity and molar activity. These initial findings warrant further exploration of its potential in detecting demyelinated lesions in the brain.

## Material and Methods

### Compliance

All rodent procedures were approved by the Animal Care and Use Committee (IACUC) at the Massachusetts General Hospital. All animal studies were conducted in compliance with the ARRIVE guidelines (Animal Research: Reporting in Vivo Experiments) for reporting animal experiments.

### Chemistry

#### General

All chemicals were ordered from commercial suppliers and used without further purification. The [^18^F]fluoride was obtained in the chemical form of [^18^F]fluoride ion ([^18^F]F^-^) via the ^18^O(p, n)^18^F nuclear reaction through bombardment of enriched [^18^O]water with 16 MeV protons at 65 µA for 6 min using a PETtrace cyclotron from GE Healthcare (GEHC) at Massachusetts General Hospital. The typical amount of [^18^F]F^-^ produced in the cyclotron target was approximately 18.5 GBq (500 mCi).

Semipreparative HPLC separations were performed by using the HPLC system connected to the TRACERlab Fx2N synthesizer (GEHC) using a Sykam S1122 Solvent Delivery System. The system was equipped with a Waters XBridge C18 Prep column (5 µm, 10 × 250 mm) and a UV detector set at 220 nm The mobile phase consisted of 20 mM sodium phosphate buffer (pH 7.4) (90%) and ethanol (10%), running at a flow rate of 4 mL/min.

The purity of the reaction crude and final compounds was analyzed using a Thermo Scientific Dionex Ultimate 3000 UHPLC system equipped with a diode array detector and a radiation detector (Model 105S, Carrol & Ramsey Instruments). Separation was carried out using an analytical Waters XBridge C18 column (5 µm, 4.6 × 150 mm) operating at a flow rate of 1 mL/min monitoring at 254 nm. The isocratic method employed a mobile phase consisting of 10 mM sodium phosphate buffer (pH 7.4) (90%) and ethanol (10%). The identity of the compound was confirmed by co-injection with a nonradioactive reference standard.

#### Azeotropic drying of the [^18^F]fluoride

The [^18^F]F^-^ was first trapped on a Sep-Pak Light Accell Plus QMA carbonate Cartridge (Waters) pre-conditioned with 5 mL 50 mM potassium bicarbonate (KHCO_3_) and 10 mL Ultrapure water. Next, the concentrated [^18^F]F^-^ was eluted into the reaction vessel using one of the following: 400 µL 50% MeCN in 0.075 M tetrabutylammounium bicarbonate (TABHCO_3_, 15 µmol) aqueous solution (ABX advanced biochemical compounds GmbH, Germany), 1.2 mL 50% MeCN in 2.5 mg/mL KHCO_3_ (15 µmol) aqueous solution or 1.2 mL 50% MeCN in tetraethylammonium bicarbonate (TEA-HCO_3_, 3 mg, 15 µmol) aqueous solution. The aqueous solution was azeotropically dried under vacuum and nitrogen flow within 14 min at 85 °C. Two aliquots of MeCN (2×500 µL) were added during the drying procedure.

#### Preparation of the precursor 1 (3-methyl-4-nitro-5-(4,4,5,5-tetramethyl-1,3,2-dioxaborolan-2-yl)pyridine)

Modifying a known procedure^24^. 3-bromo-5-methyl-4-nitropyridinemethyl (228 mg, 1.05 mmol, 1.05 equiv.), B_2_pin_2_ (254 mg, 1.0 mmol, 1.0 equiv.), Pd(dppf)Cl_2_ (146 mg, 0.2 mmol, 0.20 equiv.) and KOAc (294 mg, 3.0 mmol, 3.0 equiv.) were suspended in anhydrous and degassed 1,4-dioxane (8 mL). The mixture was stirred for 16 h at 80 °C. The mixture was allowed to cool to room temperature, diluted with CH_2_Cl_2_ and filtered through Celite. All volatiles were removed in vacuo and the residue was purified by flash column chromatography on silica (hexanes/EtOAc 67:33 to 0:100) yielding desired product in 33% yield.

#### Cu-mediated radiofluorination

A solution of Cu(OTf)_2_(py)_4_ (11 mg, 16 µmol), boronic ester **1** (3 mg, 11 µmol) in 200 µL of DMF was mixed with 100 µL of [^18^F]TEAF solution in DMF and heated up to 110 °C for 20 min. The reaction crude was diluted with ultrapure water and analyzed with radioHPLC.

#### Halogen exchange reactions

A solution of 3-bromo/chloro-5-methyl-4-nitropyridine 1-oxide (**4**, 4 mg) in 200 µL of DMSO was mixed with 100 µL of [^18^F]TBAF solution in DMSO. After stirring at room temperature for 15 min, the reaction crude was diluted with ultrapure water and analyzed with radioHPLC.

#### Radiosynthesis of [^18^F]5Me3F4AP with TRACERlab Fx2N synthesizer (GEHC)

A solution of non-radioactive 3-fluoro-5-methyl-4-nitropyridine 1-oxide (0.2 or 2 mg) in 300 µL of MeCN was added to the azeotropically dried [^18^F]F^-^. After stirring at room temperature for 1 min, the reaction was quenched by adding 3 µL of AcOH in 400 µL of MeCN solution. The reaction mixture was then dried under a nitrogen stream for 4 min.

The reaction mixture was re-dissolved in 0.8 mL of MeOH and transferred to a 4 mL V-vial charged with Pd/C (2 mg or 10 mg), a stir bar and a vent needle through a Sep-Pak Light Alumina N Cartridge (Waters). Complete transfer of the precursor was achieved by rinsing with an additional 0.8 mL of MeOH. A balloon filled with hydrogen was then installed with a needle deep into the solution, bubbling while reaction was heated to 80 °C for 10 min. The reaction mixture was then filtrated with a vent filter (Millex-FG, 0.2 µm filter unit) and rinsed with 0.8 mL of MeOH. The filtrate was transferred to Fx2N reactor vessel and dried under vacuum at room temperature for 8 min until the pressure of the reaction vessel dropped below 1 kPa. The crude was re-dissolved in water and purified by semiprep HPLC using the previously described method, with a retention time (t_R_) of approximately 30 min. The collected solution was quality checked by analytical HPLC and then diluted with saline or use as is for the animal studies.

### In vivo studies

#### Imaging Studies and Measurement of Brain SUV ex vivo

Naïve C59BI/6J mice (n = 4; male, 6– 8-week-old, Jackson Laboratory) were imaged on a Sedecal SuperArgus PET/CT scanner. Mice were injected with the radioligand (approximately 3.7 MBq in 200 µL solution) via a tail vein catheter and scanned for 60 mins in dynamic PET list mode (energy window: 250-700 KeV) under anesthesia (1.5% isoflurane with an oxygen flow of 2.0 L/min), followed by a CT scan (X Rays: 350 µA of current and 45kV of voltage). After completing the scan, the animals were euthanized, their blood was collected by cardiac puncture and their brains were harvested. Blood and brain samples were weighed, and the radioactive concentrations were measured using a single well gamma counter.

#### Imaging analysis

The images were reconstructed using a 2D ordered subset expectation maximization (OSEM) iterative reconstruction algorithm with 2 iterations and 16 subsets, including corrections for random coincident events and attenuation (using its own CT). Dynamic images were reconstructed with the following time frames 8 × 15 s, 6 × 30 s, 5 × 60 s, 4 × 300 s and remaining were 600 s frames. The obtained DICOM files were analyzed with AMIDE to generate time-activity curves with the whole brain as the region of interest.

## Abbreviations

4AP: 4-aminopyridine
BBB: blood-brain barrier
CT: computed tomography
CYP2E1: cytochrome P450 family 2 subfamily E member 1
3F4AP: 3-fluoro-4-amino-pyridine
FDA: food and drug administration
GEHC: General Electronic Healthcare
IACUC: Institutional animal care and use committee
K222: kryptofix 222
3Me4AP: 3-methyl-4-aminopyridine
5Me3F4AP: 5-methyl-3-fluoro-4-aminopyridine
MA: molar activity
MS: multiple sclerosis
PET: positron emission tomography
K_2_CO_3_: potassium carbonate
KF: potassium fluoride
K^+^: potassium ion
RCY: radiochemical yield
SNAr: nucleophilic aromatic substitution
SUV: standard uptake value
TACs: time-activity curves
TBAHCO_3_: tetrabutyl ammonium bicarbonate

## Acknowledgments

The authors also thank Kyle Stewart. David F. Lee, Jr., Dr. John A. Correia and Dr. Hamid Sabet for providing the fluorine-18 for the radiotracer synthesis.

